# Engineered protein nanocages for concurrent RNA and protein packaging *in vivo*

**DOI:** 10.1101/2022.06.16.496435

**Authors:** Seokmu Kwon, Tobias W. Giessen

## Abstract

Protein nanocages have emerged as an important engineering platform for biotechnological and biomedical applications. Among naturally occurring protein cages, encapsulin nanocompartments have recently gained prominence due to their favorable physico-chemical properties, ease of shell modification, and highly efficient and selective intrinsic protein packaging capabilities. Here, we expand encapsulin function by designing and characterizing encapsulins for concurrent RNA and protein encapsulation *in vivo*. Our strategy is based on modifying encapsulin shells with nucleic acid binding peptides without disrupting the native protein packaging mechanism. We show that our engineered encapsulins reliably self-assemble *in* vivo, are capable of efficient size-selective *in vivo* RNA packaging, can simultaneously load multiple functional RNAs, and can be used for concurrent *in vivo* packaging of RNA and protein. Our engineered encapsulation platform has potential for co-delivery of therapeutic RNAs and proteins to elicit synergistic effects, and as a modular tool for other biotechnological applications.

## Introduction

Over the past decade, protein nanocages have gained much attention for various biotechnological and biomedical applications due to their unique and desirable properties.^[1–3]^ Their biological origin makes them inherently biocompatible and allows facile genetic functionalization, while their defined shell-like structure enables the creation of multifunctional and atomically defined nano-devices by modifying both their inner and outer surfaces. Further, established recombinant protein production strategies make protein-based nanostructures simple to produce, purify, and scale.

Based on these favorable properties, significant effort has been dedicated towards engineering protein nanocages like bacterial microcompartments (BMCs),^[4,5]^ lumazine synthase,^[6,7]^ ferritin,^[8,9]^ virus-like particles (VLPs),^[10,11]^ and computationally designed protein shells.^[12]^ For example, BMCs have been engineered as catalytic nanoreactors^[13–15]^ and molecular scaffolds,^[16]^ while lumazine synthase has been utilized for molecular display,^[17–19]^ RNA packaging and delivery,^[20,21]^ and enzyme encapsulation.^[22,23]^ Ferritins and VLPs have long been used for *in vitro* bionanotechnology, biomaterials research, and therapeutics delivery.^[8–11]^ More recently, designed protein assemblies have similarly been explored for related applications.^[20,24–26]^

Among naturally occurring protein nanocages, encapsulins have emerged as an alternative and attractive engineering platform for applications in medicine, catalysis, and nanotechnology.^[27–33]^ Encapsulins are self-assembling icosahedral protein compartments composed of a single type of shell protomer possessing the HK97 phage-like fold.^[34–36]^ They can assemble into T1 (60 subunits, ca. 24 nm),^[36–39]^ T3 (180 subunits, ca. 32 nm),^[40,41]^ and T4 (240 subunits, ca. 42 nm)^[42]^ shells and are widely distributed throughout both the bacterial and archaeal domains.^[43,44]^ Encapsulins have been proposed to play diverse roles in cellular metabolism including iron storage,^[42]^ redox stress resistance,^[45]^ and sulfur metabolism.^[38]^ Their key feature is the ability to selectively encapsulate dedicated cargo proteins *in vivo*.^[36]^ All native cargo proteins contain N- or C-terminal domains^[38]^ or targeting peptides (TPs) necessary for efficient cargo loading during shell self-assembly.^[46,47]^ This feature – a dedicated and modular protein loading mechanism – has been widely utilized to package non-native cargo proteins into the encapsulin shell via simple genetic fusion of TPs to proteins of interest.^[48–50]^

Engineered encapsulins have shown potential as nanoreactors,^[50,51]^ drug delivery systems,^[52]^ imaging agents,^[53–55]^ and immunotherapies.^[30,56,57]^ Straightforward genetic and chemical shell modification allows small molecule conjugation,^[33]^ peptide loop insertion,^[58]^ pore modification,^[50,59,60]^ and fusion of protein domains to the N- and C-terminus of the encapsulin protomer.^[50,52,56,61]^ Recently, engineered encapsulins capable of triggered reversible disassembly, enabling *in vitro* cargo loading and stimulus-responsive cargo release, have also been reported.^[62]^

A topic of particular current interest is the selective packaging and delivery of nucleic acids inside protein-based cages.^[1,63]^ Engineering such systems has shed light on the evolution and function of viruses and allowed the creation of non-viral systems mimicking select virus characteristics.^[21,64–66]^ The ability to encapsulate nucleic acids *in vivo* may provide novel approaches for RNA regulation and cytosolic sampling^[21]^ with broad implications for RNA biology.^[67–70]^ RNA- and DNA-based therapeutics have tremendous clinical potential.^[71,72]^ However, their broad application has been hampered by poor pharmacokinetic properties,^[73]^ difficulty in overcoming cell membranes,^[74]^ susceptibility to nucleases, inherent immunogenicity, and rapid clearance from the body.^[75,76]^ These challenges could be overcome by engineering efficient nucleic acid delivery systems, with many different approaches and materials already having been employed towards achieving this goal.^[73,77–79]^ Due to their desirable properties and engineerability, protein cages in general and encapsulins in particular, represent a promising alternative strategy for nucleic acid packaging and delivery. In addition, expanding encapsulin function towards nucleic acid encapsulation would allow for the concurrent sequestration and co-localization of proteins and nucleic acids. This may enable the future co-delivery of two types of functional macromolecules acting either in an orthogonal or synergistic manner.

Here, we engineer and characterize encapsulins as novel nano-encapsulation platforms for simultaneous RNA and protein packaging *in vivo*, laying the foundation for their future use as targeted co-delivery systems.

## Results and Discussion

### Design and initial characterization of an encapsulin-based *in vivo* RNA encapsulation system

We set out to design robust and modular encapsulin-based nanocages for the *in vivo* sequestration of RNA, without disturbing their native protein loading capabilities. Further, the three naturally occurring assembly states of encapsulins – T1, T3, and T4 – were exploited to design a range of RNA packaging nanocages with different dimensions. This strategy allows the exploration of the influence of luminal volume and charge on *in vivo* RNA and protein loading. In particular, the three established encapsulin systems from *Thermotoga maritima* (TmT1), *Myxococcus xanthus* (MxT3), and *Quasibacillus thermotolerans* (QtT4) were chosen as engineering scaffolds (Figure 1A). Although the Mx encapsulin primarily forms T3 shells, a recent study showed that in the absence of native protein cargo, about 36% of shells assemble into small T1-sized encapsulins with a diameter of 18 nm (MxT1).^[40]^ This feature was used to additionally explore the RNA packaging capacity of MxT1.

**Figure 1.**
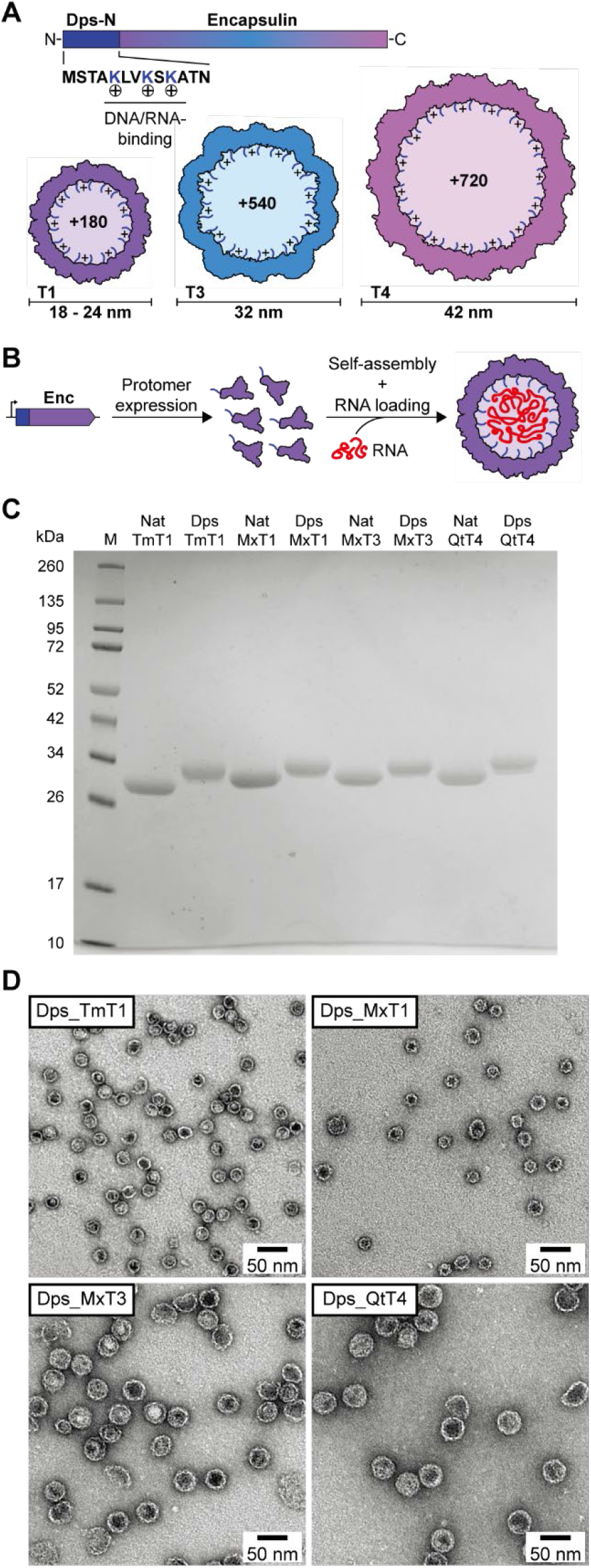
Design and initial characterization of the encapsulin-based *in vivo* RNA encapsulation system. A) Schematic of the engineered encapsulin protomer sequence with the N-terminally fused nucleic acid-binding peptide Dps-N. The three different sizes (T1, T3, and T4) of engineered encapsulins (Dps_Encs) created in this study are shown highlighting the additional luminal positive charges introduced by Dps-N. T: triangulation number (T-number). B) Schematic outlining the principle of *in vivo* nucleic acid encapsulation by Dps_Encs. Enc: encapsulin. C) SDS-PAGE analysis and comparison of purified native (Nat) and engineered (Dps) encapsulins used in this study. Encapsulins are labeled by their organism of origin and T-number. Tm: *Thermotoga maritima*, Mx: *Myxococcus xanthus*, Qt: *Quasibacillus thermotolerans*. D) Negative-stain TEM micrographs of all four purified Dps_Encs used in this study.

To imbue encapsulins with the ability to bind and encapsulate nucleic acid, we genetically fused the *Escherichia coli* Dps-N peptide (MSTAKLVKSKATN) – originating from the DNA-binding protein from starved cells (Dps)^[80]^ – to the N-terminus of the encapsulin protomer via a flexible 6 residue linker (GGSGGS) yielding our Dps_Enc fusion constructs. Dps-N consists of the 13 N-terminal residues of Dps and includes 3 positively charged lysines (Figure 1A). It is able to bind to both DNA and RNA, likely via the electrostatic interaction of the positively charged lysine residues with the negatively charged DNA/RNA phosphate backbone. Dps-N was specifically chosen for our fusion constructs due to its broad specificity and prior successful use as a nucleic acid-binding peptide.^[81]^ In assembled Tm, Mx, and Qt encapsulins, the N-termini of all protomers are pointed towards the shell interior. Therefore, in engineered Dps_Encs, 3 additional positive charges per protomer will be introduced to the encapsulin lumen, resulting in overall charge increases of +180 (T1), +540 (T3), and +720 (T4) for our fusion constructs (Figure 1A). This increased positive charge of the shell interior will drive the encapsulation of RNA during shell self-assembly (Figure 1B). We envisioned that Dps_Encs would allow *in vivo* packaging of native or overexpressed RNAs whilst minimizing concurrent DNA packaging. This is due to the relatively small size of encapsulins and the fact that generally, no DNA molecules small enough to be encapsulated inside encapsulin shells are present inside cells. Further, the broad specificity of Dps-N could in the future also be used for the flexible *in vitro* loading of variable nucleic acids, both RNA and DNA.

Dps_Encs and unmodified native Tm, Mx, and Qt controls (Nat_Encs) were produced in *E. coli* and purified through a combination of polyethylene glycol (PEG) precipitation, ion exchange chromatography (IEC), and size-exclusion chromatography (SEC). SDS-PAGE analysis of purified Dps_Encs and Nat_Encs was used to confirm sample homogeneity (Figure 1C). Further analyses using negative-stain transmission electron microscopy (TEM), dynamic light scattering (DLS), and analytical SEC indicated that all Dps_Encs formed stable shells with similar size and appearance compared to the corresponding Nat_Encs (Figure 1D, Figure S1). These results confirm the feasibility of our novel Dps-N fusion designs and highlight the ease of luminal encapsulin shell modification without disturbing shell self-assembly.

### *In vivo* nucleic acid encapsulation and resistance towards nuclease digestion

After confirming the proper assembly of all Dps_Enc designs, we next focused on their nucleic acid encapsulation capacity and ability to protect encapsulated nucleic acid from nuclease digestion (Figure 2A). Native agarose gel electrophoresis with both protein and nucleic acid staining showed that all purified Dps_Encs contained significantly more nucleic acid than the respective Nat_Enc controls (Figure 2B, middle). To exclude non-specific nucleic acid binding to the outside of encapsulin shells, Benzonase treatments and IEC were carried out during all purifications. These results confirm that Dps-N fusion does indeed confer nucleic acid encapsulation capacity to all of our engineered encapsulin shells, further supported by direct A260/A280 measurements of purified samples, and increased A260/A280 signal observed during SEC (Figure S1).

**Figure 2.**
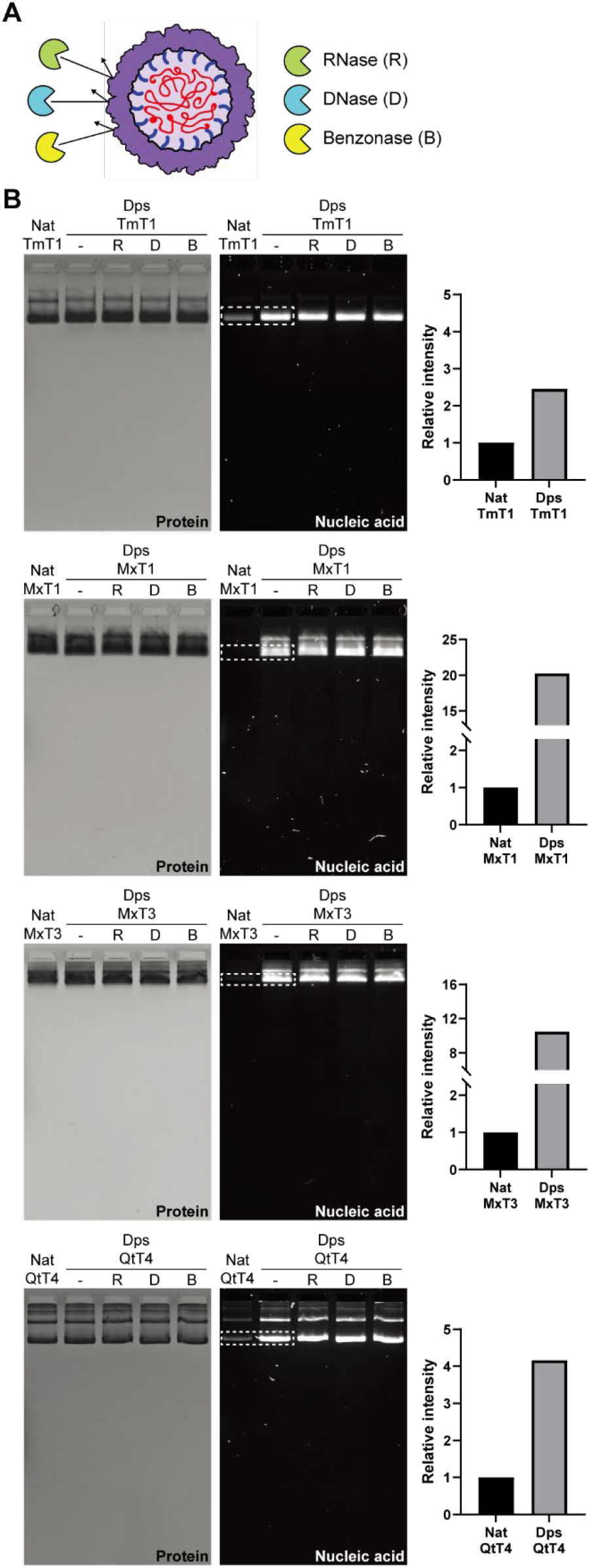
*In vivo* nucleic acid encapsulation inside Dps_Encs and resistance towards nuclease digestion. A) Schematic of Dps_Encs protecting *in vivo* encapsulated nucleic acids from nucleases. B) Native agarose gel electrophoresis of Nat_Encs and Dps_Encs before and after nuclease treatment. Gels were stained with Coomassie Blue to visualize protein (left) or GelRed to visualize nucleic acids (middle). On the right, the relative intensity of nucleic acid bands (dotted box) normalized by protein amount is shown, highlighting the encapsulation capacity difference between Nat_Encs and Dps_Encs.

Dps_MxT1 and Dps_MxT3 showed the highest relative nucleic acid packaging capacity with 20- and 11-fold increases in signal when compared to their native forms (Figure 2B, right). Dps_TmT1 and Dps_QtT4 yielded more moderate signal increases of 2.4- and 4-fold, respectively. These discrepancies in increased nucleic acid loading capacity over native encapsulins are partially caused by the larger background signals observed for Nat_TmT1 and Nat_QtT4. For both Nat_Encs and Dps_Encs, several minor higher molecular weight bands could be observed. We confirmed via tryptic digest and mass spectrometry that all observed bands represent the respective Nat_Encs or Dps_Encs (Figure S2). Higher molecular weight bands are likely a result of partial aggregation of encapsulin shells during gel electrophoresis.

Next, purified Dps_Encs were treated with DNase, RNase, or Benzonase. No reduction in the intensity of nucleic acid bands was observed for any of our fusion constructs, confirming that the encapsulin shell can effectively protect encapsulated nucleic acid from nuclease digestion (Figure 2B, middle). The protective role of encapsulin shells is likely due to the physical sequestration of nucleic acid inside a protein barrier. Encapsulin shells possess small pores at the 5-, 3-, and 2-fold symmetry axes with diameters ranging from 2 - 7 Å,^[40,42,59]^ which is too small to allow nuclease access to the shell interior. Further, encapsulin shells are generally very stable and once formed can only be disassembled under harsh non-physiological conditions, thus making them excellent containers for protecting labile nucleic acids.

### Analysis of encapsulated nucleic acid content and size-selective RNA packaging

To identify the type and size distribution of encapsulated nucleic acid, we first extracted the total nucleic acid contents from purified Dps_Encs and subjected them to differential nuclease treatment using DNase, RNase, or Benzonase (Figure 3).

**Figure 3.**
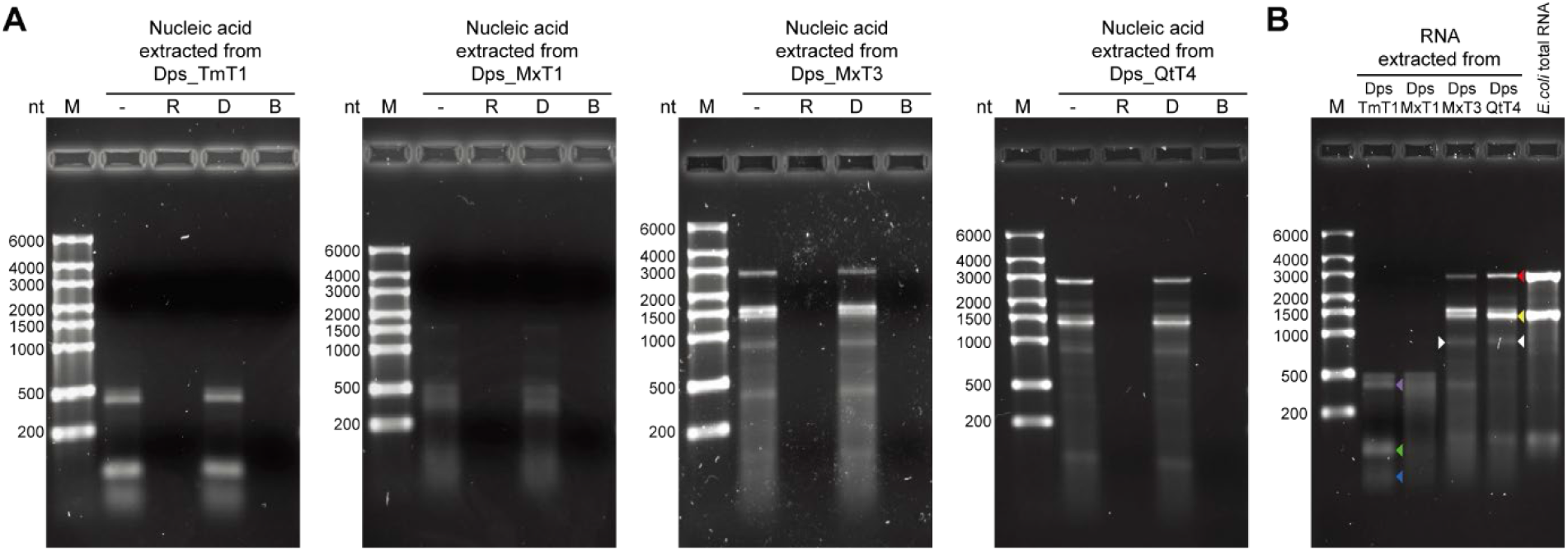
Analysis of encapsulated nucleic acid content and size-selective RNA packaging of Dps_Encs. A) Native agarose gel electrophoresis of nucleic acid extracted from purified Dps_Encs treated with either RNase (R), DNase (D), or Benzonase (B). Only nucleic acid treated with DNase retained its integrity whereas RNase and Benzonase treatment led to complete digestion. B) Native agarose gel electrophoresis of RNA extracted from Dps_Encs along with *E*.*coli* total RNA showing the size selectivity of differently sized Dps_Encs. A differential band for Dps_MxT3 and Dps_QtT4 - absent in *E. coli* total RNA or T1 Dps_Encs - is highlighted by white arrows and putatively represents the respective Dps_Enc mRNA. Colored arrows indicate: 23S rRNA (red), 16S rRNA (yellow), tmRNA (purple), 5S rRNA (green), and tRNA (blue).

Exposure to DNase had no effect on any of the extracted nucleic acid samples, while RNase and Benzonase treatment resulted in complete digestion (Figure 3A). This clearly indicates that nucleic acid encapsulated in Dps_Encs is exclusively RNA, thus confirming our initial design for *in vivo* RNA packaging based on the fact that generally, only DNA molecules too large to be encapsulated inside encapsulin shells exist in cells, specifically chromosomes and plasmids.

We further compared the size distribution of RNA extracted from purified Dps_Encs. RNA encapsulation capacity was found to be size-selective and proportional to shell size, with smaller encapsulins showing a lower upper size limit for RNA compared to larger shells (Figure 3B). Specifically, Dps_TmT1 and Dps_MxT1 exhibited a maximum size of encapsulated RNA of ∼500 nt, whereas the larger Dps_MxT3 and Dps_QtT4 shells were able to encapsulate RNA of up to ∼3,000 nt in length. Comparison of extracted RNA with *E. coli* total RNA indicated that for Dps_TmT1 and Dps_MxT1, tRNAs (blue arrow), 5S rRNA (green arrow), and tmRNA (purple arrow) likely made up a substantial part of sequestered RNA, while for Dps_MxT3 and Dps_QtT4, 16S (yellow arrow) and 23S (red arrow) rRNA were found to be the main RNA species (Figure 3B). This result is in accordance with rRNA and tRNA generally representing the majority of available RNA inside cells. Because the molecular size of RNA depends on its ability to form secondary structures, with many functional RNAs even able to adopt stable and compact 3D folds, RNAs larger than the observed size limits could potentially be encapsulated in Dps_Encs as well. Further, differential bands of ∼1000 nt in length, absent in *E*.*coli* total RNA and T1 Dps_Encs, were observed for Dps_MxT3 and Dps_QtT4 (Figure 3B, white arrows). Given that the mRNA size of Dps_MxT3 and Dps_QtT4 transcripts is ∼1000 nt, and that they would have been overexpressed, these differential bands likely represent Dps_MxT3 and Dps_QtT4 mRNA. Overall, these results indicate that Dps_Encs are able to encapsulate RNA in a size-selective manner with RNA abundance also playing an important role in determining encapsulation efficiency.

### Analysis of RNA packaging capacity

To determine the relative RNA packaging capacities of our Dps_Encs per encapsulin shell, we performed native polyacrylamide gel electrophoresis (PAGE), loading the same normalized amount of encapsulin shells per lane for all Dps_Encs (Figure 4A). We found that RNA packaging capacity per shell increases with shell diameter (Figure 4B). Larger Dps_Encs have larger volumes for RNA packaging and contain more Dps-N-fused protomers which results in an increased number of positive luminal charges. This indicates that RNA encapsulation capacity for Dps_Encs correlates with the overall number of charges and the available shell volume, rather than approximate luminal charge density which would be maximal for Dps_MxT1 (Figure S3).

**Figure 4.**
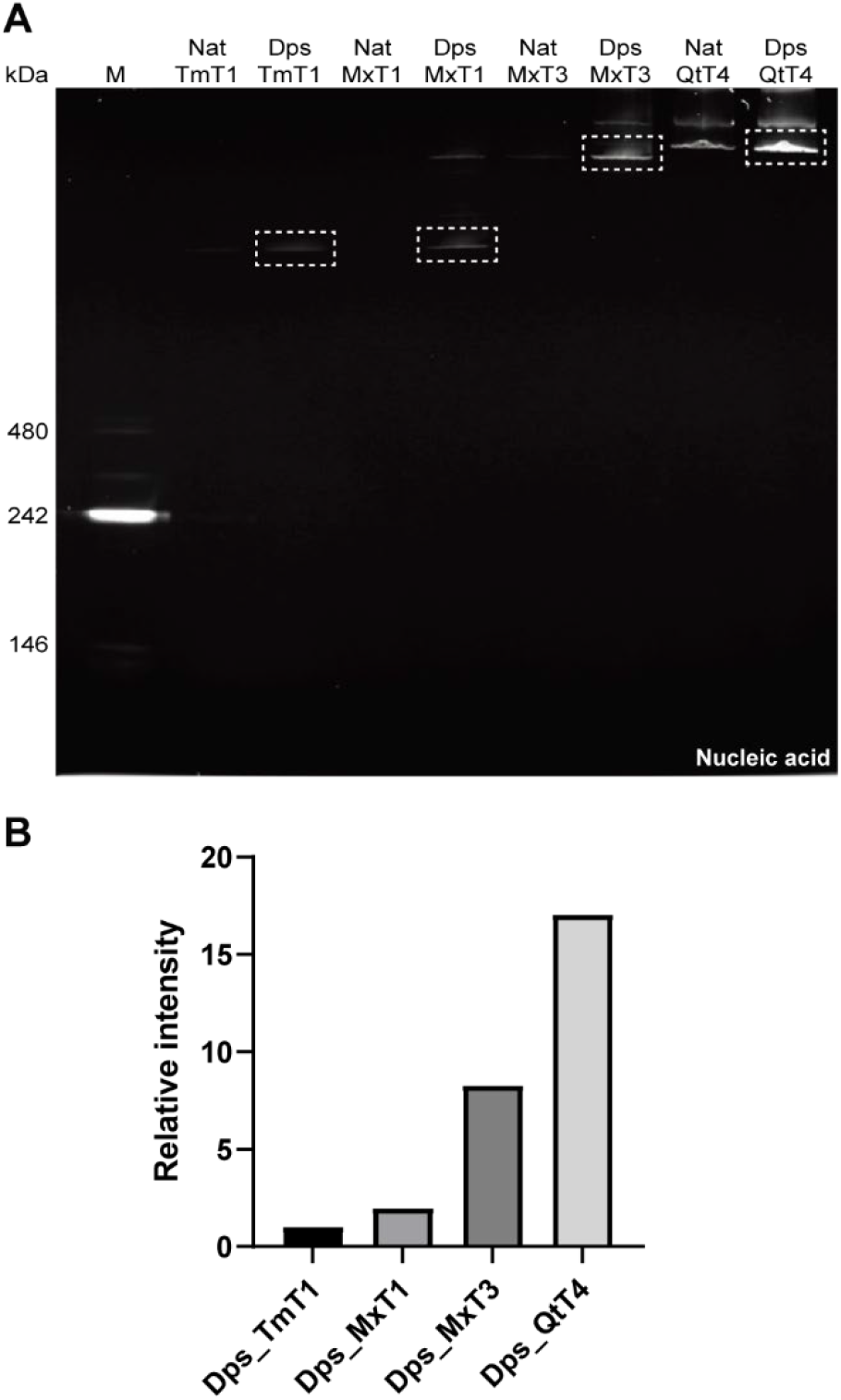
RNA packaging capacity analysis of differently sized Nat_Encs and Dps_Encs. A) Native PAGE gel analysis of Nat_Encs and Dps_Encs stained with GelRed to visualize RNA where equivalent amounts of Dps_Encs are loaded (normalized to the number of Dps_Enc shells per lane) across all lanes for comparative analysis. B) Relative intensity of RNA bands (dotted boxes in A) showing that per Dps_Enc shell, RNA packaging capacity increases with the shell size of Dps_Encs.

### Simultaneous *in vivo* packaging of two functional RNAs

To expand the utility of Dps_Encs, we sought to test if multiple non-endogenous functional RNAs could be co-packaged at the same time and be protected from nucleases. We utilized the split fluorogenic aptamer Split_Broccoli (SB)^[82]^ and co-expressed its two RNAs – Top (97 nt) and Bottom (153 nt) – together with Dps_MxT3 (Figure 5A, Figure S4).

**Figure 5.**
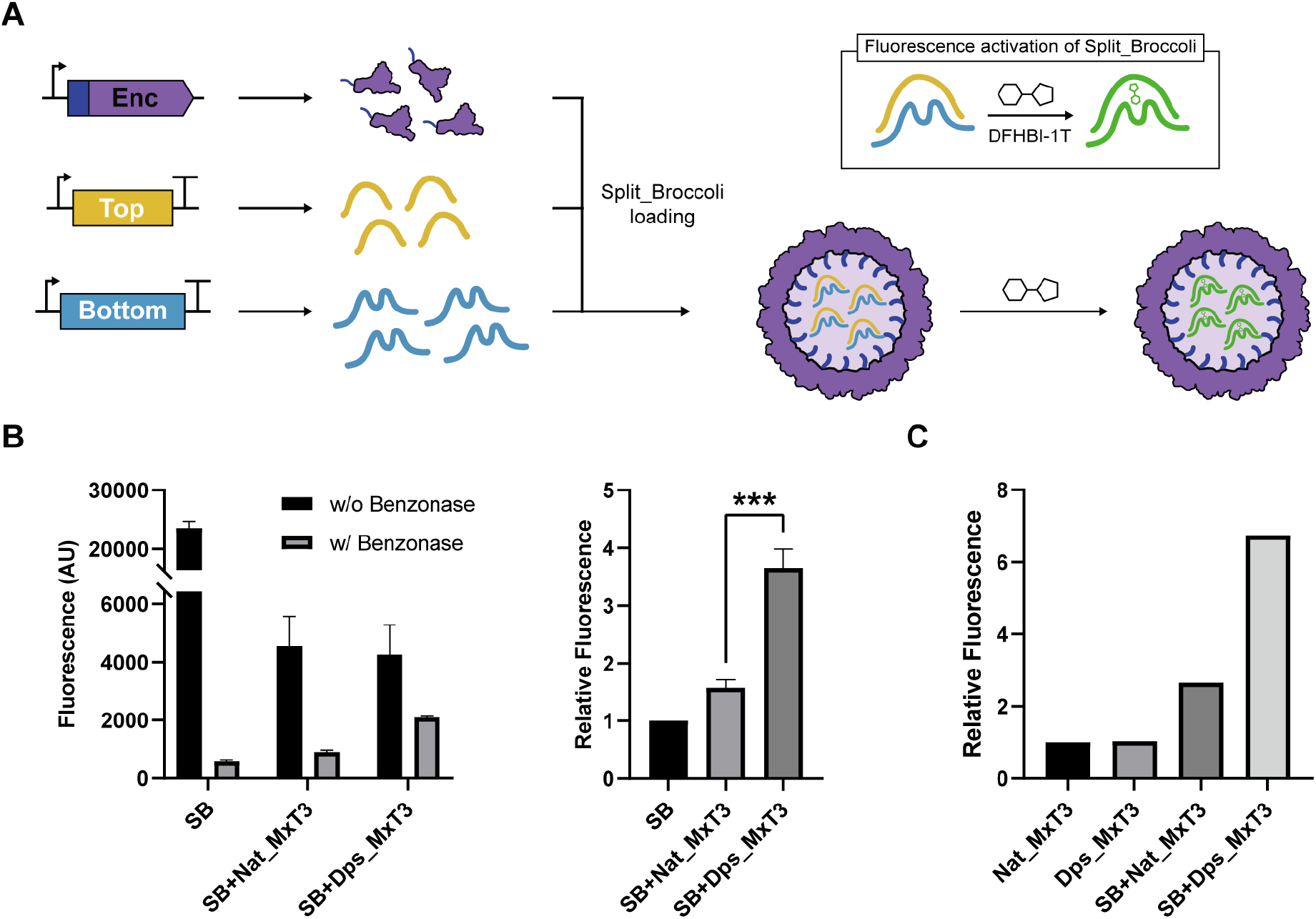
Simultaneous *in vivo* packaging of two functional Split_Brocolli (SB) RNAs using Dps_MxT3. A) Schematic outlining the expression strategy and fluorescence activation of the SB aptamer (box). After *in vivo* packaging of both SB RNAs, SB fluorescence can be activated by the addition of the small fluorogenic molecule DFHBI-1T. SB is composed of the Top (yellow) and Bottom (blue) RNAs. B) Fluorescence measurements (left) of DFHBI-1T-supplemented cell lysates - from cells expressing SB, SB + Nat_MxT3, or SB + Dps_MxT3 - with or without prior Benzonase treatment. Relative fluorescence (right) of Benzonase-treated cell lysates normalized by the fluorescence of Benzonase-treated SB cell lysate. Data are shown as mean values while error bars represent standard deviations from three independent experiments. (***P = 0.0005, two-sided unpaired t-test) C) Relative fluorescence of purified and DFHBI-1T-supplemented Nat_MxT3 and Dps_MxT3 that were expressed in cells with or without concurrent SB expression. Relative fluorescence was normalized based on the fluorescence of the Nat_MxT3 control.

Dps_MxT3 was used due to its overall favorable performance, combining a high upper size limit for RNA, low background, and high loading capacity. In initial experiments, Benzonase was added to cleared cell lysates – from cells expressing SB alone, SB + Nat_MxT3, or SB + Dps_MxT3 – to remove free SB, highlight the protective role of encapsulin shells, and allow the detection of encapsulated SB via the addition of the small molecule SB binding partner DFHBI-1T, yielding a fluorescence readout (Figure 5B). The highest SB fluorescence signal was observed for SB + Dps_MxT3 (Figure 5B), indicating that Dps_MxT3 successfully packaged both SB RNAs, protected them from nuclease digestion, and allowed access of the small molecule DFHBI-1T to the shell interior. To confirm these experiments, Nat_MxT3 and Dps_MxT3 were purified alone or from cells co-expressing SB, followed by incubation with DFHBI-1T. Again, substantially higher SB fluorescence signal was observed for Dps_MxT3, confirming our initial results (Figure 5C).

### Concurrent RNA and protein packaging *in vivo*

To design a system for the simultaneous *in vivo* packaging of both RNA and protein, we sought to combine the newly engineered ability of our Dps_Encs to encapsulate RNA with encapsulins’ native capacity for specific protein encapsulation. As a proof of concept, the Mx targeting peptide (MxTP, PEKRLTVGSLRR) with flexible 6 residue linker (GGSGGS) was genetically fused to the C-terminus of eGFP and cloned immediately upstream of the Dps_MxT3 gene for co-expression (Figure 6A). SDS-PAGE analysis of purified Dps_MxT3 confirmed the successful *in vivo* loading and co-purification of MxTP-tagged eGFP (Figure 6B). Negative stain TEM analysis further confirmed that eGFP-loaded Dps_MxT3 particles still formed homogeneous shells, very similar in size and appearance to Nat_MxT3 (Figure 6C, Figure S1). To test concurrent loading of both eGFP and RNA, native PAGE analysis was performed on purified eGFP-loaded Dps_MxT3 shells. Compared to Nat_MxT3, eGFP-loaded Dps_MxT3 exhibited substantially higher RNA signal intensity and eGFP fluorescence of the high molecular weight encapsulin band. Co-elution of RNA and eGFP signals confirms successful co-packaging of RNA and a specific heterologously expressed protein *in vivo*. This novel ability could be useful for future biomedical delivery applications where therapeutic effects may be potentiated by the synergistic action of co-delivered functional RNAs and proteins.

**Figure 6.**
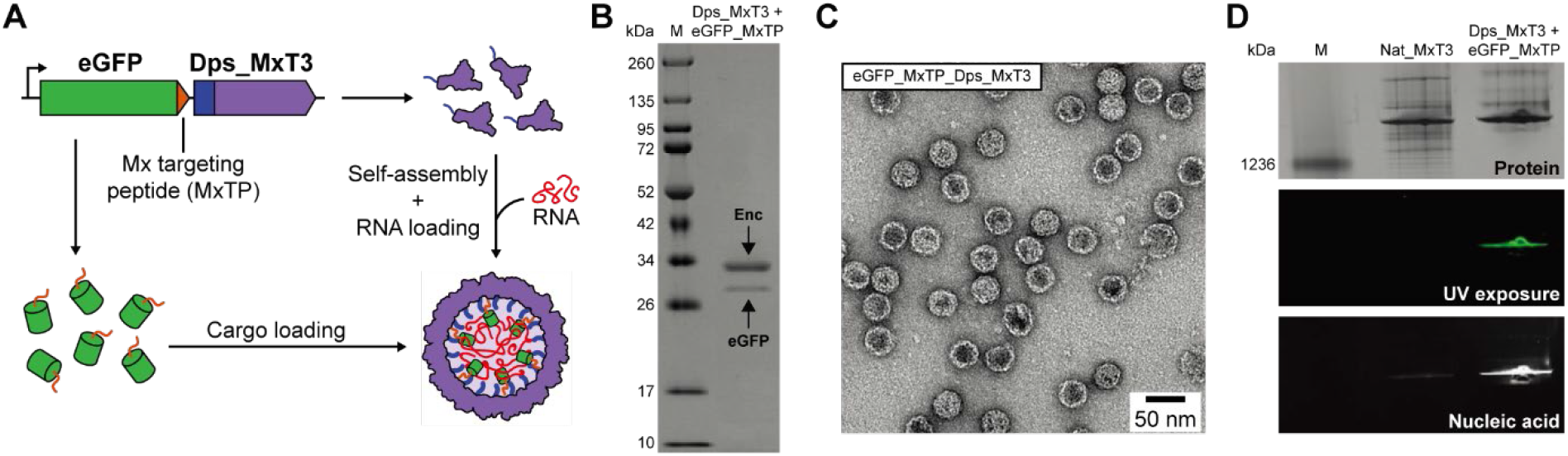
Concurrent *in vivo* RNA and protein packaging using Dps_MxT3. A) Schematic of Dps_MxT3 concurrently packaging RNA and eGFP. RNA is packaged through interaction with the fused Dps-N peptide while eGFP is loaded through specific targeting peptide (MxTP) interaction. B) SDS-PAGE analysis of purified eGFP- and RNA-loaded Dps_MxT3. C) Negative-stain TEM micrograph of purified eGFP- and RNA-loaded Dps_MxT3. D) Native PAGE analysis showing Nat_MxT3 and eGFP- and RNA-loaded Dps_MxT3 via Coomassie blue staining (top), UV exposure for eGFP fluorescence analysis (middle), and GelRed staining for RNA detection (bottom).

## Conclusion

Protein nanocage engineering has the potential to significantly contribute innovative solutions to challenging problems across various fields, including catalysis, nanotechnology, and medicine. In particular, innovative nanocage designs can yield high-performing tools for biomedical delivery applications. Delivery modalities based on protein cages have so far focused on safely and efficiently transporting protein- or RNA-based therapeutics into target cells.^[20,68,83,84]^ However, the possibility of combining two types of specific therapeutic macromolecules – protein and RNA – in a single nanocage design, has barely been explored.^[85]^ One advantage of co-delivering, in principle, any combination of therapeutic RNAs and proteins of interest to cells include the potential for targeting multiple intracellular target classes, e.g., mRNA and protein, at the same time. For example, combining siRNA and antibodies against the same target in a single nanocage-based delivery vehicle may lead to improved suppression due to dual action at both the mRNA and protein level. A second advantage of co-packaging RNA and protein in a single nanocage is that it leads to ensured co-delivery to each target cell, whereas concurrent administration of the same therapeutics via separate delivery methods has the potential to lead to heterogeneous populations of singly and doubly targeted cells.^[86]^

To address the challenge of co-packaging multiple types of functional macromolecules in a single nanocage design, we have developed encapsulin-based protein cages, called Dps_Encs, capable of concurrently packaging functional RNAs and specific proteins of interest *in vivo*. Importantly, all Dps_Encs efficiently protected encapsulated RNA from nuclease digest which is one of the main challenges of RNA-based therapeutics (Figure 3).^[87]^ Four different sizes of Dps_Encs were created, namely, MxT1 (18 nm, luminal volume: ∼905 nm^3^), TmT1 (24 nm, ∼3,054 nm^3^), MxT3 (32 nm, ∼9,203 nm^3^), and QtT4 (42 nm, ∼24,429 nm^3^). The available luminal volume spans more than an order of magnitude and correlates well with the observed RNA loading capacity per shell (Figure 4). We further showed that Dps_MxT3 is capable of co-localizing and protecting two functional RNAs, the split aptamer Split_Broccoli, and importantly, that the SB binding partner DFHBI-1T can access the shell interior, likely via the 5-, 3-, or 2-fold pores natively present in MxT3 (Figure 5). Finally, we showed that Dps_MxT3 retained the ability to specifically sequester a TP-tagged co-expressed cargo protein whilst simultaneously packaging RNA (Figure 6).

While most protein cage systems to date rely on *in vitro* packaging of cargo,^[88–91]^ requiring separate purification as well as disassembly and reassembly steps, our encapsulin-based Dps_Encs can co-package RNA and specific proteins *in vivo* in a single step. This *in situ* assembly of functional nanocages simplifies purification and avoids non-physiological *in vitro* conditions, often necessary for disassembly and cargo loading of other protein nanocages.^[88–91]^ One challenge of *in vivo* cargo loading is the potential co-packaging of unwanted molecules, including endogenous RNA and protein. However, the intrinsic specificity of encapsulins for packaging co-expressed TP-tagged proteins has been extensively used to assemble highly homogeneous cargo-loaded cages with minimal non-specific loading.^[46–50]^ After purification, *in vivo* eGFP-loaded Dps_MxT3 showed minimal background of non-TP-tagged proteins (Figure 6B). In future encapsulin-based nanocage designs, Dps-N could easily be replaced with RNA-binding peptides or domains that bind RNA in a sequence-specific manner.^[92–94]^ Functional RNAs could then be tagged with this packaging RNA sequence likely resulting in sequence-selective *in vivo* RNA loading. Another potential advantage of nanocages that can be loaded *in vivo* is their potential use in living therapeutics.^[95]^ In living therapeutics, engineered bacteria are used as a drug delivery modality to reach a target site of interest. Once at the target, bioactive molecules can be continuously produced locally by the bacteria, leading to increased therapeutic effects with minimal systemic side effects.^[96,97]^ Thus, nanocage systems that do not require *in vitro* assembly, could be locally assembled *in vivo* and released.^[96,98]^ Finally, Dps_Encs could themselves be imbued with cell targeting capabilities through the genetic fusion of cell penetrating peptides or similar targeting systems, to the encapsulin C-terminus exposed on the shell exterior.^[33,52]^

In sum, the Dps_Encs developed in this study lay the foundation for using encapsulins for the co-delivery of therapeutic RNA and protein with the ultimate goal of eliciting homogeneous synergistic effects at a single cell level. This work further highlights the versatility of encapsulins as modular and robust tools with broad potential applicability for different biomedical and biotechnological applications.

## Acknowledgements

We gratefully acknowledge funding from the NIH (R35GM133325) (TWG) and a Korean Government Scholarship (SK).

## Conflict of Interest

The authors declare no conflict of interest.

### Experimental Section Molecular biology and cloning

All constructs used in this study, except Dps-N-fused encapsulins (Dps_Encs), were ordered from Integrated DNA Technologies (IDT) as *E. coli* codon-optimized gBlocks (Table S1). Genes for Dps_Encs were obtained through overhang PCR using the native encapsulin genes as templates, adding Dps-N in the process. All genes except Split_Broccoli (SB) were cloned into the pETDuet-1 vector while SB was inserted into pCDFDuet-1 using Gibson Assembly. *E. coli* BL21 (DE3) cells were transformed with the assembled plasmids via electroporation and were confirmed through Sanger sequencing (Eurofins Scientific).

### Protein expression and purification

All expression experiments were carried out using lysogeny broth (LB) medium supplemented with the appropriate selection marker (100 mg/mL ampicillin (pETDuet-1), 50 mg/mL spectinomycin (pCDFDuet-1), or both). 500 mL of fresh LB medium were inoculated 1:100 using a 5 mL overnight culture, grown at 37°C to OD600 of 0.4-0.5, and then induced with 0.1 mM IPTG. After induction, cultures were grown at 30°C overnight for ca. 18 h and harvested via centrifugation (8,000 g, 10 min, 4°C). The resulting cell pellets were frozen and stored at - 20°C until further use.

Frozen cell pellets were resuspended in 5 mL/g (wet cell mass) of Tris buffer (20 mM Tris, 150 mM NaCl, pH 7.5). Lysis components [lysozyme (0.5 mg/mL), Benzonase® nuclease (25 units/mL), MgCl_2_ (1.5 mM), SIGMAFAST EDTA-free protease inhibitor cocktail (one tablet per 100 mL)] were added and cells were incubated on ice for 15 min. Samples were then sonicated at 55% amplitude and a pulse time of 10 s on and 20 s off for 5 min total (Model 120 Sonic Dismembrator, Fisher Scientific). After sonication, samples were clarified by centrifugation (10,000 g, 15 min, 4°C). To the supernatant, NaCl and PEG-8000 were added to a final concentration of 0.5 M and 10%, respectively, and incubated on ice for 40 min, followed by centrifugation (8000 g, 10 min, 4°C). The supernatant was removed, and the pellet was resuspended in 3 mL Tris buffer (pH 7.5) and filtered using a 0.2 μm syringe filter.

The filtered sample was subjected to SEC using a Sephacryl S-500 16/60 column and Tris buffer (pH 7.5) at a flow rate of 1 mL/min. Fractions were evaluated using SDS-PAGE and encapsulin-containing fractions were combined, concentrated, and dialyzed using Amicon filter units (100 kDa MWCO) and Tris buffer without NaCl (20 mM Tris, pH 7.5). The low salt sample was then loaded on a HiPrep DEAE FF 16/10 Ion Exchange column at a flow rate of 3 mL/min to remove nucleic acid contamination. Encapsulin-containing fractions were concentrated, centrifuged (10,000 g, 10 minutes, 4°C), and then subjected to SEC using a Superose 6 10/300 GL column and Tris buffer (pH 7.5) at a flow rate of 0.5 mL/min. Purified proteins were stored in Tris buffer (pH 7.5) at 4°C until further use.

### Transmission electron microscopy

Encapsulin samples for negative-stain TEM were diluted to 0.15 mg/mL in Tris buffer (pH 7.5). Gold grids (200-mesh coated with Formvar-carbon film, EMS) were made hydrophilic by glow discharge at 5 mA for 60 s (easiGlow, PELCO). 4 μL of sample was added to the grid and incubated for 1 min, wicked with filter paper, and washed with 0.75% uranyl formate before staining with 0.75% uranyl formate for 1 min. Stain was removed using filter paper and the grid was dried for at least 20 min before imaging. TEM micrographs were captured using a Morgagni transmission electron microscope at 100 keV at the University of Michigan Life Sciences Institute.

### Dynamic light scattering (DLS) analysis

All sizing and polydispersity measurements were carried out on an Uncle instrument (Unchained Labs) at 15°C in triplicate. All encapsulin samples were adjusted to 0.5 mg/mL of monomer using Tris buffer (pH 7.5), centrifuged (10,000 g, 10 minutes, 4°C), and then immediately analyzed via DLS.

### RNA extraction

RNA was extracted from purified Dps_Enc samples via Phenol-Chloroform extraction followed by ethanol precipitation. Phenol:chloroform:isoamyl alcohol (25:24:1, pH 8) was used for Phenol-Chloroform extraction and after ethanol precipitation, the desalted nucleic acid extracts were dissolved in TEN buffer (Tris 10 mM, EDTA 1 mM, pH 8) and stored at -80°C. *E. coli* total RNA was purchased from ThermoFisher Scientific (AM7940). Quantification of RNA was carried out using a Nanodrop Spectrophotometer from ThermoFisher Scientific, Inc. (USA).

### Nuclease challenge of extracted RNA and RNA-loaded Dps_Encs

DNase (ThermoFisher Scientific, EN0521), RNase (ThermoFisher Scientific, EN0531), and Benzonase (Sigma Aldrich, E8263) were used for nuclease challenge experiments of extracted RNA and RNA-loaded Dps_Encs. For all nuclease incubation experiments, 1 μL (1.5 μL) of DNase, RNase, and Benzonase were added to μL (13.5 μL) of extracted RNA (RNA-loaded Dps_Encs) samples (final concentration: 10 U/mL, 5 U/mL, and 25 U/mL, respectively) followed by 30 min incubation at 37°C.

### Native gel electrophoresis

#### Native agarose gel electrophoresis

3% native agarose gels were used to determine the nucleic acid encapsulation capacity of Dps_Encs and to demonstrate the nuclease resistance of Dps_Encs shell. The amount of Dps_Enc loaded per lane was adjusted for each Dps_Enc encapsulin so as to easily visualize nucleic acid signal after GelRed staining, while corresponding Nat_Encs were loaded at equal amounts for direct comparison. Per lane, 15 μL of sample were loaded with an additional 2 μL of 70% (v/v) aqueous glycerol. Gel electrophoresis was carried out using 1X TAE buffer at a constant voltage of 90 V for 35 min. Gels were first stained with GelRed to visualize nucleic acids and then stained with Coomassie Blue to visualize proteins. Nucleic acid encapsulation capacity of Nat_Encs and Dps_Encs was compared via gel densitometry, first, normalizing the intensity of nucleic acid bands by their corresponding protein band (N/P). Then, N/P values of Dps_Encs were normalized by N/P values of the corresponding Nat_Encs for comparison. Band intensities of nucleic acid and protein were measured using Fiji/ImageJ v2.1.0/1.53c. 2% native agarose gels were used for nucleic acid extracted from purified Dps_Encs. The extracted nucleic acid was incubated with nucleases and loaded on the gels along with undigested nucleic acid for comparison. Per lane, 10 μL of sample were loaded with an additional10 μL of 2x RNA loading buffer. Gel electrophoresis was carried out in 1X TAE buffer at a constant voltage of 125 V for 25-30 min. The gel was stained with GelRed to visualize nucleic acids.

#### Native polyacrylamide gel electrophoresis

All native polyacrylamide gel electrophoresis analyses were conducted in an Invitrogen XCell SureLock using NativePAGE™ 3 to 12% bis-tris mini protein gels with 1X NativePAGE™ Anode Buffer and 1X NativePAGE™ Cathode Buffer. 850 fmol of encapsulin shells were loaded per lane to maintain equivalent amounts of shells across all lanes for comparative analysis. Native PAGE gels were run at a constant voltage of 150 V for 1 hour followed by an additional 1 hour run at 250 V at 4°C. Gels were then stained, first with GelRed for nucleic acid visualization, then with Coomassie Blue for protein detection. For eGFP_MxTP_Dps_MxT3, the gel was first exposed to UV light for eGFP visualization before staining with GelRed and Coomassie Blue.

To quantify and compare the amount of RNA loaded in each Dps_Enc encapsulin, gel densitometry of GelRed-stained gels was carried out using Fiji/ImageJ v2.1.0/1.53c. Pixel intensities of bands were background subtracted yielding final overall intensities per band for comparisons.

### Split_Broccoli (SB) fluorescence experiments

50 mL of fresh LB medium containing appropriate antibiotic(s) were inoculated using an overnight 1 mL culture of either SB-, SB+Nat_MxT3-, or SB+Dps_MxT3-expressing cells. Cultures were grown at 37°C to an OD600 of 0.4-0.5, then induced with 0.2 mM IPTG and further grown for 5 hours at 30°C. As a control, 50 mL of *E*.*coli* BL21 (DE3) without transformed plasmids were similarly grown in LB at 30°C for 5 hours. Harvested cells were resuspended in 5 mL of Tris buffer (pH 7.5) and sonicated at 55% amplitude and a pulse time of 10 s on and 20 s off for 3 min 30 s total. Lysates were clarified by centrifugation (10,000 g, 15 min, 4°C) and supernatants were filtered using 0.2 μm syringe filters. For each sample, two 100 μL aliquots were prepared. To one of the two aliquots of each sample, 14.7 μL of DFHBI-1T (final concentration: 1 mM) was added and incubated at 37°C for 40 min, followed by fluorescence measurements using a Synergy H1 plate reader configured with filter sets for green fluorescence (λ_ex_ = 472 nm, λ_em_ = 507 nm). For the other aliquot, 1 μL of MgCl_2_ (final concentration: 1.5 mM) and 1 μL of Benzonase (250 units) were added and incubated overnight at room temperature. The following day, 15 μL of DFHBI-1T (final concentration: 1 mM) was added and incubated at 37°C for 40 min, followed by fluorescence analysis. For fluorescence measurements, 25 μL of each sample were loaded per well in triplicate into a black-flat bottom 384-well plate. Background fluorescence from the control *E*.*coli* BL21(DE3) sample without plasmid was subtracted from all samples yielding final fluorescence intensities.

SB fluorescence was also measured using purified Nat_MxT3, Dps_MxT3, SB+Nat_MxT3, and SB+Dps_MxT3 samples. To 75 μL of each sample containing 5 pmol of capsid, 2 μL of DFHBI-1T (final concentration: 200 uM) was added and incubated at 37°C for 40 min, followed by fluorescence analysis as described above. As background, 75 μL of Tris buffer (pH 7.5) was used and subtracted from the fluorescence signal of each sample.

## References

[1] T. G. W. Edwardson, M. D. Levasseur, S. Tetter, A. Steinauer, M. Hori, D. Hilvert, Chemical Reviews 2022, DOI 10.1021/acs.chemrev.1c00877.

[2] T. W. Giessen, P. A. Silver, Journal of Molecular Biology 2016, 428, 916–927.

[3] S. Bhaskar, S. Lim, NPG Asia Materials 2017, 9, 1–18.

[4] S. Planamente, S. Frank, Biochemical Society Transactions 2019, 47, 765–777.

[5] J. S. Plegaria, C. A. Kerfeld, Current Opinion in Biotechnology 2018, 51, 1–7.

[6] Y. Azuma, T. G. W. Edwardson, D. Hilvert, Chemical Society Reviews 2018, 47, 3543–3557.

[7] Y. Wei, P. Kumar, N. Wahome, N. J. Mantis, C. R. Middaugh, Journal of Pharmaceutical Sciences 2018, 107, 2283–2296.

[8] G. Jutz, P. van Rijn, B. Santos Miranda, A. Böker, Chemical Reviews 2015, 115, 1653–1701.

[9] N. Song, J. Zhang, J. Zhai, J. Hong, C. Yuan, M. Liang, Accounts of Chemical Research 2021, 54, 3313–3325.

[10] Fei. Hill, Brett.; Zak, Andrew.; Khera, Eshita. Wen, Current Protein & Peptide Science 2018, 19, 112–127.

[11] S. Nooraei, H. Bahrulolum, Z. S. Hoseini, C. Katalani, A. Hajizade, A. J. Easton, G. Ahmadian, Journal of Nanobiotechnology 2021, 19, 1–27.

[12] I. Stupka, J. G. Heddle, Current Opinion in Structural Biology 2020, 64, 66–73.

[13] A. D. Lawrence, S. Frank, S. Newnham, M. J. Lee, I. R. Brown, W. F. Xue, M. L. Rowe, D. P. Mulvihill, M. B. Prentice, M. J. Howard, M. J. Warren, ACS Synthetic Biology 2014, 3, 454–465.

[14] M. Liang, S. Frank, H. Lünsdorf, M. J. Warren, M. B. Prentice, Biotechnology Journal 2017, 12, 1600415.

[15] H. Kirst, B. H. Ferlez, S. N. Lindner, C. A. R. Cotton, A. Bar-Even, C. A. Kerfeld, Proc Natl Acad Sci U S A 2022, 119, 1–10.

[16] G. Zhang, T. Johnston, M. B. Quin, C. Schmidt-Dannert, ACS Synthetic Biology 2019, 8, 1867–1876.

[17] Y. Wei, N. Wahome, G. VanSlyke, N. Whitaker, P. Kumar, M. L. Barta, W. L. Picking, D. B. Volkin, N. J. Mantis, C. R. Middaugh, Protein Science 2017, 26, 2059–2072.

[18] Y. Komatsu, N. Terasaka, K. Sakai, E. Mihara, R. Wakabayashi, K. Matsumoto, D. Hilvert, J. Takagi, H. Suga, iScience 2021, 24, 103302.

[19] M. D. Levasseur, S. Mantri, T. Hayashi, M. Reichenbach, S. Hehn, Y. Waeckerle-Men, P. Johansen, D. Hilvert, ACS Chemical Biology 2021, 16, 838–843.

[20] T. G. W. Edwardson, T. Mori, D. Hilvert, J Am Chem Soc 2018, 140, 10439–10442.

[21] Y. Azuma, T. G. W. Edwardson, N. Terasaka, D. ilvert, J Am Chem Soc 2018, 140, 566–569.

[22] Donald. Azuma, Yusuke.; Hilvert, in Protein Scaffolds, 2018, pp. 39–55.

[23] Y. Azuma, D. L. V. Bader, D. Hilvert, J Am Chem Soc 2018, 140, 860–863.

[24] T. G. W. Edwardson, M. D. Levasseur, D. Hilvert, Chimia (Aarau) 2021, 75, 323–328.

[25] K. A. Cannon, R. U. Park, S. E. Boyken, U. Nattermann, S. Yi, D. Baker, N. P. King, T. O. Yeates, Protein Science 2020, 29, 919–929.

[26] T. E. Harcus, M. Gluckman, H. Pontzer, D. A. Raichlen, F. W. Marlowe, W. R. Siegfried, I. A. W. Macdonald, J. Call, J. Fischer, K. F. Stryjewski, S. Quader, M. D. Sorenson, N. Boogert, N. Davies, T. Flower, G. Jamie, R. Magrath, D. Rendall, G. Ruxton, M. Sorensen, B. Wood, C. David, Science (1979) 2016, 353, 389–395.

[27] T. W. Giessen, Current Opinion in Chemical Biology 2016, 34, 1–10.

[28] T. W. Giessen, P. A. Silver, Current Opinion in Biotechnology 2017, 46, 42–50.

[29] A. Groaz, H. Moghimianavval, F. Tavella, T. W. Giessen, A. G. Vecchiarelli, Q. Yang, A. P. Liu, Wiley Interdisciplinary Reviews: Nanomedicine and Nanobiotechnology 2021, 13, 1–27.

[30] P. Lagoutte, C. Mignon, G. Stadthagen, S. Potisopon, S. Donnat, J. Mast, A. Lugari, B. Werle, Vaccine 2018, 36, 3622–3628.

[31] T. W. Giessen, P. A. Silver, ACS Synthetic Biology 2016, 5, 1497–1504.

[32] T. H. Lee, T. S. Carpenter, P. D’haeseleer, D. F. Savage, M. C. Yung, Biotechnology and Bioengineering 2020, 117, 603–613.

[33] H. Moon, J. Lee, J. Min, S. Kang, Biomacromolecules 2014, 15, 3794–3801.

[34] T. W. Giessen, Annual Review of Biochemistry 2022, 91, 1–28.

[35] M. P. Andreas, T. W. Giessen, Nature Communications 2021, 12, 1–16.

[36] M. Sutter, D. Boehringer, S. Gutmann, S. Günther, D. Prangishvili, M. J. Loessner, K. O. Stetter, E. Weber-Ban, N. Ban, Nature Structural and Molecular Biology 2008, 15, 939–947.

[37] J. Ross, Z. McIver, T. Lambert, C. Piergentili, J. E. Bird, K. J. Gallagher, F. L. Cruickshank, P. James, E. Zarazúa-Arvizu, L. E. Horsfall, K. J. Waldron, M. D. Wilson, C. Logan Mackay, A. Baslé, D. J. Clarke, J. Marles-Wright, Science Advances 2022, 8, 1–12.

[38] R. J. Nichols, B. Lafrance, N. R. Phillips, D. R. Radford, L. M. Oltrogge, L. E. Valentin-Alvarado, A. J. Bischoff, E. Nogales, D. F. Savage, Elife 2021, 10, 1–22.

[39] N. Lončar, H. J. Rozeboom, L. E. Franken, M. C. A. Stuart, M. W. Fraaije, Biochemical and Biophysical Research Communications 2020, 529, 548–553.

[40] E. Eren, B. Wang, D. C. Winkler, N. R. Watts, A. C. Steven, P. T. Wingfield, Structure 2022, 30, 551–563.

[41] F. Akita, K. T. Chong, H. Tanaka, E. Yamashita, N. Miyazaki, Y. Nakaishi, M. Suzuki, K. Namba, Y. Ono, T. Tsukihara, A. Nakagawa, Journal of Molecular Biology 2007, 368, 1469– 1483.

[42] T. W. Giessen, B. J. Orlando, A. A. Verdegaal, M. G. Chambers, J. Gardener, D. C. Bell, G. Birrane, M. Liao, P. A. Silver, Elife 2019, 8, 1–23.

[43] T. W. Giessen, P. A. Silver, Nature Microbiology 2017, 2, 1–11.

[44] J. C. Tracey, M. Coronado, T. W. Giessen, M. C. Y. Lau, P. A. Silver, B. B. Ward, Scientific Reports 2019, 9, 1–11.

[45] K. A. Lien, R. J. Nichols, C. Cassidy-Amstutz, K. Dinshaw, M. Knight, R. Singh, L. D. Eltis, D. F. Savage, S. A. Stanley, bioRxiv 2020, 2020.08.31.276014.

[46] W. J. Altenburg, N. Rollins, P. A. Silver, T. W. Giessen, Scientific Reports 2021, 11, 1–9.

[47] C. Cassidy-Amstutz, L. Oltrogge, C. C. Going, A. Lee, P. Teng, D. Quintanilla, A. East-Seletsky, E. R. Williams, D. F. Savage, Biochemistry 2016, 55, 3461–3468.

[48] Y. H. Lau, T. W. Giessen, W. J. Altenburg, P. A. ilver, Nature Communications 2018, 9, 1–7.

[49] F. Sigmund, C. Massner, P. Erdmann, A. Stelzl, H. Rolbieski, M. Desai, S. Bricault, T. P. Wörner, J. Snijder, A. Geerlof, H. Fuchs, M. H. de Angelis, A. J. R. Heck, A. Jasanoff, V. Ntziachristos, J. Plitzko, G. G. Westmeyer, Nature Communications 2018, 9, 1–14.

[50] M. C. Jenkins, S. Lutz, ACS Synthetic Biology 2021, 10, 857–869.

[51] P. Lohner, M. Zmyslia, J. Thurn, J. K. Pape, R. Gerasimaitė, J. Keller-Findeisen, S. Groeer, B. Deuringer, R. Süss, A. Walther, S. W. Hell, G. Lukinavičius, T. Hugel, C. Jessen-Trefzer, Angewandte Chemie - International Edition 2021, 60, 23835–23841.

[52] A. van de Steen, R. Khalife, N. Colant, H. Mustafa Khan, M. Deveikis, S. Charalambous, C. M. Robinson, R. Dabas, S. Esteban Serna, D. A. Catana, K. Pildish, V. Kalinovskiy, K. Gustafsson, S. Frank, Synthetic and Systems Biotechnology 2021, 6, 231–241.

[53] A. N. Gabashvili, S. S. Vodopyanov, N. S. Chmelyuk, V. A. Sarkisova, K. A. Fedotov, M. v. Efremova, M. A. Abakumov, International Journal of Molecular Sciences 2021, 22, 12275.

[54] M. v. Efremova, S. V. Bodea, F. Sigmund, A. Semkina, G. G. Westmeyer, M. A. Abakumov, Pharmaceutics 2021, 13, 397.

[55] Y. Zhang, X. Wang, C. Chu, Z. Zhou, B. Chen, X. Pang, G. Lin, H. Lin, Y. Guo, E. Ren, P. Lv, Y. Shi, Q. Zheng, X. Yan, X. Chen, G. Liu, Nature Communications 2020, 11, 1–11.

[56] M. Kanekiyo, W. Bu, M. G. Joyce, G. Meng, J. R. R. Whittle, U. Baxa, T. Yamamoto, S. Narpala, J. P. Todd, S. S. Rao, A. B. McDermott, R. A. Koup, M. G. Rossmann, J. R. Mascola, B. S. Graham, J. I. Cohen, G. J. Nabel, Cell 2015, 162, 1090–1100.

[57] B. Choi, H. Moon, S. J. Hong, C. Shin, Y. Do, S. Ryu, S. Kang, ACS Nano 2016, 10, 7339– 7350.

[58] S. Michel-Souzy, N. M. Hamelmann, S. Zarzuela-Pura, J. M. J. Paulusse, J. J. L. M. Cornelissen, Biomacromolecules 2021, 22, 5234–5242.

[59] L. S. R. Adamson, N. Tasneem, M. P. Andreas, W. Close, E. N. Jenner, T. N. Szyszka, R. Young, L. C. Cheah, A. Norman, H. I. MacDermott-Opeskin, M. L. O’Mara, F. Sainsbury, T. W. Giessen, Y. H. Lau, Science Advances 2022, 8, 1–13.

[60] E. M. Williams, S. M. Jung, J. L. Coffman, S. Lutz, ACS Synthetic Biology 2018, 7, 2514–2517.

[61] H. Choi, S. Eom, H. U. Kim, Y. Bae, H. S. Jung, S. Kang, Biomacromolecules 2021, 22, 3028–3039.

[62] J. A. Jones, A. S. Cristie-David, M. P. Andreas, T. W. Giessen, Angewandte Chemie - International Edition 2021, 60, 25034–25041.

[63] M. Comas-Garcia, M. Colunga-Saucedo, S. Rosales-Mendoza, M. Comas-Garcia, Molecular Pharmaceutics 2020, 17, 4407–4420.

[64] N. Terasaka, Y. Azuma, D. Hilvert, Proc Natl Acad Sci U S A 2018, 115, 5432–5437.

[65] T. G. W. Edwardson, D. Hilvert, J Am Chem Soc 2019, 141, 9432–9443.

[66] S. Tetter, N. Terasaka, A. Steinauer, R. J. Bingham, S. Clark, A. J. P. Scott, N. Patel, M. Leibundgut, E. Wroblewski, N. Ban, P. G. Stockley, R. Twarock, D. Hilvert, Science (1979) 2021, 372, 1220–1224.

[67] P. Y. Fang, L. M. G. Ramos, S. Y. Holguin, C. Hsiao, J. C. Bowman, H. W. Yang, L. D. Williams, Nucleic Acids Research 2017, 45, 3519–3527.

[68] K. M. Choi, K. Kim, I. C. Kwon, I. S. Kim, H. J. Ahn, Molecular Pharmaceutics 2013, 10, 18–25.

[69] S. Qazi, H. M. Miettinen, R. A. Wilkinson, K. McCoy, T. Douglas, B. Wiedenheft, Molecular Pharmaceutics 2016, 13, 1191–1196.

[70] L. Ponchon, M. Catala, B. Seijo, M. el Khouri, F. Dardel, S. Nonin-Lecomte, C. Tisné, Nucleic Acids Research 2013, 41, 1–13.

[71] C. I. Edvard Smith, R. Zain, Annual Review of Pharmacology and Toxicology 2019, 59, 605– 630.

[72] K. E. Lundin, O. Gissberg, C. I. E. Smith, Human Gene Therapy 2015, 26, 475–485.

[73] K. A. Whitehead, R. Langer, D. G. Anderson, Nature Reviews Drug Discovery 2009, 8, 129– 138.

[74] S. F. Dowdy, Nature Biotechnology 2017, 35, 222–229.

[75] M. P. Gantier, B. R. G. Williams, Cytokine and Growth Factor Reviews 2007, 18, 363–371.

[76] F. Iversen, C. Yang, F. Dagnæs-Hansen, D. H. Schaffert, J. Kjems, S. Gao, Theranostics 2013, 3, 201–209.

[77] N. C. Seeman, Nano Letters 2020, 20, 1477– 1478.

[78] T. Lehto, K. Ezzat, M. J. A. Wood, S. el Andaloussi, Advanced Drug Delivery Reviews 2016, 106, 172–182.

[79] Y. Ding, Z. Jiang, K. Saha, C. S. Kim, S. T. Kim, R. F. Landis, V. M. Rotello, Molecular Therapy 2014, 22, 1075–1083.

[80] L. N. Calhoun, Y. M. Kwon, Journal of Applied Microbiology 2011, 110, 375–386.

[81] C. Park, Y. Jin, Y. J. Kim, H. Jeong, B. L. Seong, Biochemical and Biophysical Research Communications 2020, 524, 484–489.

[82] K. K. Alam, K. D. Tawiah, M. F. Lichte, D. Porciani, D. H. Burke, ACS Synthetic Biology 2017, 6, 1710–1721.

[83] Y. Wang, M. Uchida, H. K. Waghwani, T. Douglas, ACS Synthetic Biology 2020, 9, 3298– 3310.

[84] S. J. Kaczmarczyk, K. Sitaraman, H. A. Young, S. H. Hughes, D. K. Chatterjee, Proc Natl Acad Sci U S A 2011, 108, 16998–17003.

[85] S. Banskota, A. Raguram, S. Suh, S. W. Du, J. R. Davis, E. H. Choi, X. Wang, S. C. Nielsen, G. A. Newby, P. B. Randolph, M. J. Osborn, K. Musunuru, K. Palczewski, D. R. Liu, Cell 2022, 185, 250-265.e16.

[86] P. M. Glassman, V. R. Muzykantov, Journal of Pharmacology and Experimental Therapeutics 2019, 370, 570–580.

[87] J. C. Kaczmarek, P. S. Kowalski, D. G. Anderson, Genome Medicine 2017, 9, 1–16.

[88] F. C. Herbert, O. R. Brohlin, T. Galbraith, C. Benjamin, C. A. Reyes, M. A. Luzuriaga, A. Shahrivarkevishahi, J. J. Gassensmith, J. J. Gassensmith, Bioconjugate Chemistry 2020, 31, 1529–1536.

[89] O. Tagit, M. v. de Ruiter, M. Brasch, Y. Ma, J. J. L. M. Cornelissen, RSC Advances 2017, 7, 38110–38118.

[90] J. Zhang, D. Cheng, J. He, J. Hong, C. Yuan, M. Liang, Nature Protocols 2021, 16, 4878–4896.

[91] I. Stupka, Y. Azuma, A. P. Biela, M. Imamura, S. Scheuring, E. Pyza, O. Woźnicka, D. P. Maskell, J. G. Heddle, Science Advances 2022, 8, DOI 10.1126/sciadv.abj9424.

[92] R. J. Austin, T. Xia, J. Ren, T. T. Takahashi, R. W. Roberts, J Am Chem Soc 2002, 124, 10966–10967.

[93] S. Hyun, K. H. Lee, A. Han, J. Yu, Nucleic Acid Therapeutics 2011, 21, 157–163.

[94] J. D. Puglisi, L. Chen, S. Blanchard, A. D. Frankel, Science 1995, 270, 1200–1203.

[95] A. Cubillos-Ruiz, T. Guo, A. Sokolovska, P. F. Miller, J. J. Collins, T. K. Lu, J. M. Lora, Nature Reviews Drug Discovery 2021, 20, 941–960.

[96] C. R. Gurbatri, I. Lia, R. Vincent, C. Coker, S. Castro, P. M. Treuting, T. E. Hinchliffe, N. Arpaia, T. Danino, Science Translational Medicine 2020, 12, 1–12.

[97] D. S. Leventhal, A. Sokolovska, N. Li, C. Plescia, S. A. Kolodziej, C. W. Gallant, R. Christmas, J. Gao, M. J. James, A. Abin-fuentes, M. Momin, C. Bergeron, A. Fisher, P. F. Miller, K. A. West, J. M. Lora, Nature Communications 2020, 11, 1–15.

[98] A. Bar-Zion, A. Nourmahnad, D. R. Mittelstein, S. Shivaei, S. Yoo, M. T. Buss, R. C. Hurt, D. Malounda, M. H. Abedi, A. Lee-Gosselin, M. B. Swift, D. Maresca, M. G. Shapiro, Nature Nanotechnology 2021, 16, 1403–1412.

