## Supplementary Information for "Engineered protein nanocages for concurrent RNA and protein packaging *in vivo*"

A

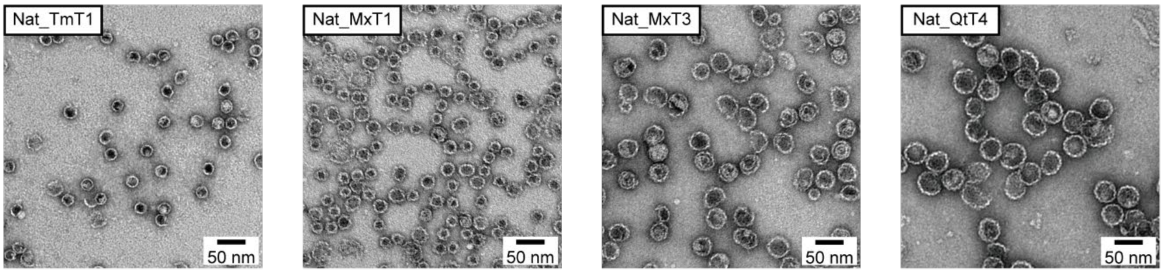

Nat_TmT1
50 nm
Nat_MxT1
50 nm
Nat_MxT3
50 nm
Nat_QtT4
50 nm


B

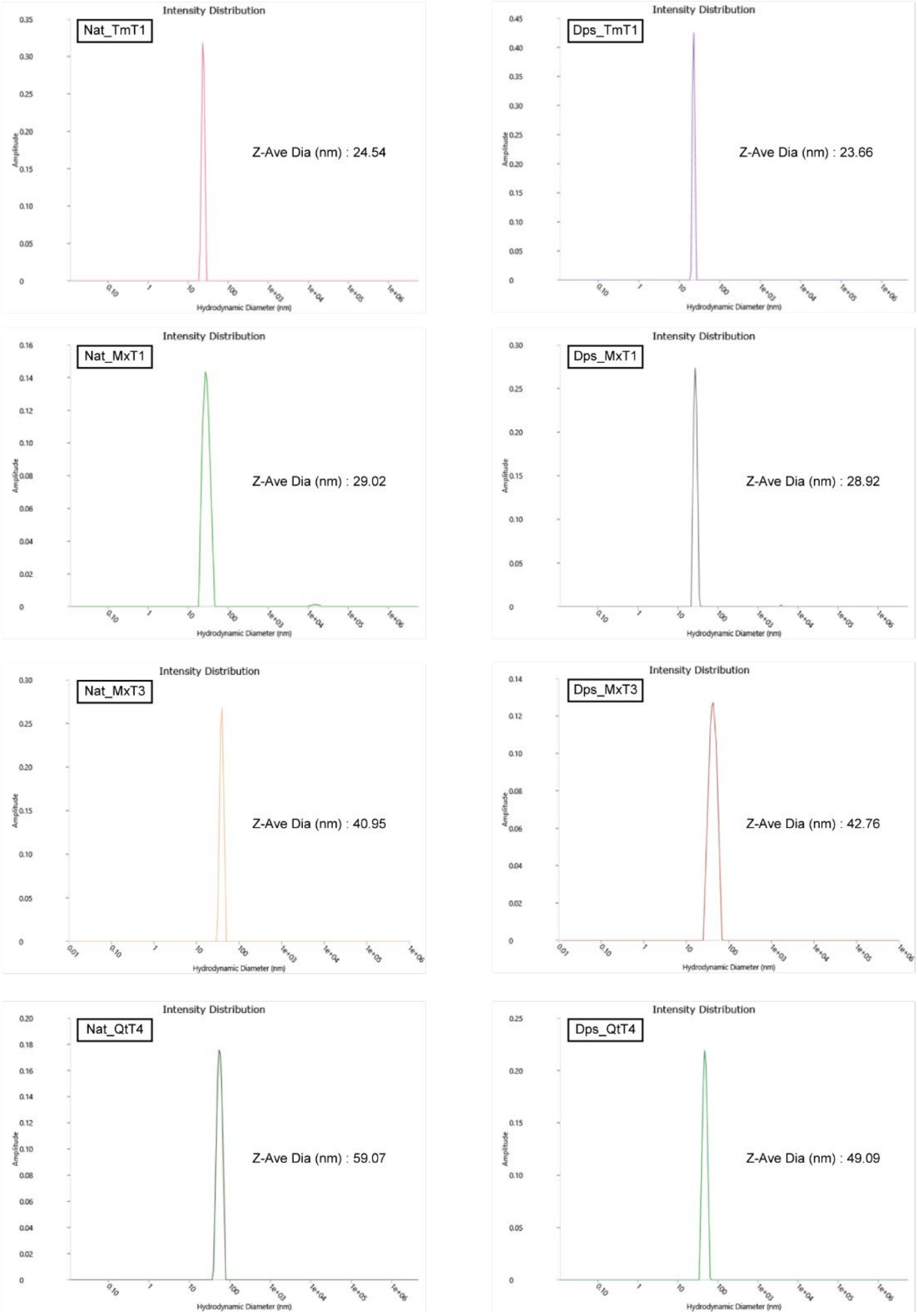

Intensity Distribution
Nat_TmT1
Z-Ave Dia (nm) : 24.54
Dps_TmT1
Z-Ave Dia (nm) : 23.66
Nat_MxT1
Z-Ave Dia (nm) : 29.02
Dps_MxT1
Z-Ave Dia (nm) : 28.92
Nat_MxT3
Z-Ave Dia (nm) : 40.95
Dps_MxT3
Z-Ave Dia (nm) : 42.76
Nat_QtT4
Z-Ave Dia (nm) : 59.07
Dps_QtT4
Z-Ave Dia (nm) : 49.09
Amplitude
Hydrodynamic Diameter (nm)

**C**

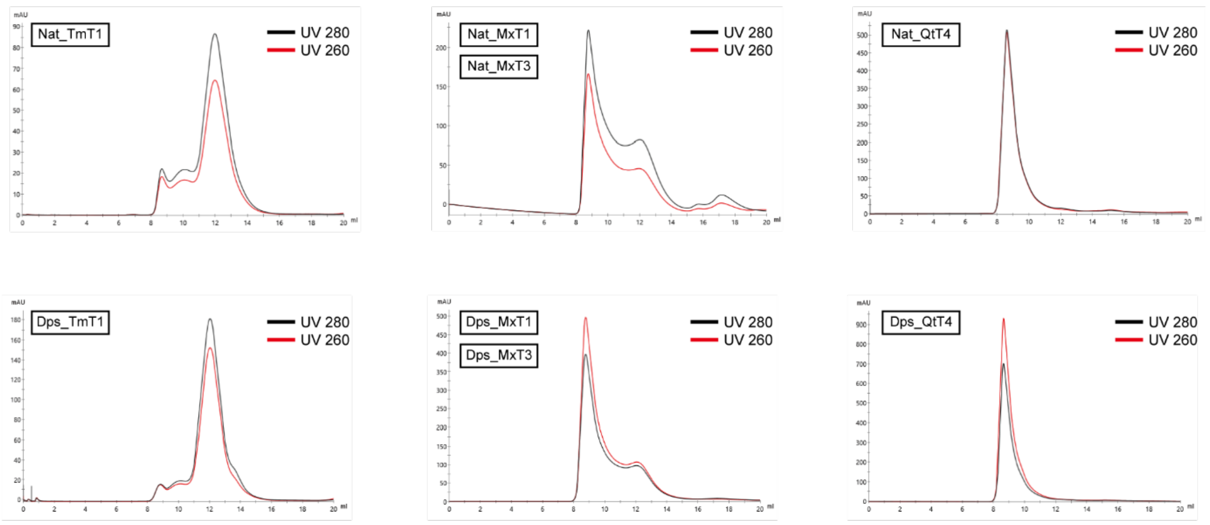


**D**

| | Collected elution volume from SEC (ml) | A260/A280 |
|---|---|---|
| Nat_TmT1 | 11 – 13 | 0.76 |
| Dps_TmT1 | 11 – 13 | 0.86 |
| Nat_MxT1 | 11 – 13 | 0.64 |
| Dps_MxT1 | 11 – 13 | 1.11 |
| Nat_MxT3 | 8 – 10 | 0.77 |
| Dps_MxT3 | 8 – 10 | 1.24 |
| Nat_QtT4 | 8 – 10 | 0.98 |
| Dps_QtT4 | 8 – 10 | 1.33 |

**Figure S1.** A) Negative-stain TEM micrographs of all four purified Nat_Encs. B) Dynamic light scattering (DLS) analysis of Nat_Encs (left) and Dps_Encs (right). Z-average diameter for each sample is shown next to the peak. C) Size-exclusion chromatography (SEC) analysis of Nat_Encs (top) and Dps_Encs (bottom). UV 280 tracking the protein signal is shown in black line and UV 260 tracking the nucleic acid signal is shown in red line. D) Table showing the collected elution volume for each Nat_Enc and Dps_Enc from the respective SEC runs (second column) and the direct A260/A280 measurement of the collected and concentrated Nat_Encs and Dps_Encs (third column).

**A**

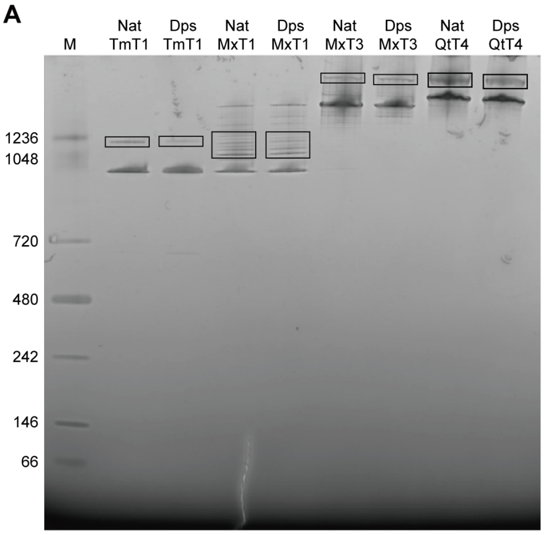


**B**

| Identified Proteins (74) | Accession Number | Molecular Weight | SpC Sample 1 |
|---|---|---|---|
| Nat_TmT1\|SequenceA\|SequenceA | Custom\|SequenceA\|Seque | 30 kDa | 42 |
| 50S ribosomal protein L20 OS=Escherichia coli (strain K12) OX=83333 GN=rplT PE=1 | sp\|P0A7L3\|RL20_ECOLI | 13 kDa | 14 |
| Elongation factor Tu 1 OS=Escherichia coli (strain K12) OX=83333 GN=tufA PE=1 SV | sp\|P0CE47\|EFTU1_ECOLI (+ | 43 kDa | 13 |
| 50S ribosomal protein L2 OS=Escherichia coli (strain K12) OX=83333 GN=rplB PE=1 | sp\|P60422\|RL2_ECOLI | 30 kDa | 12 |
| Glyceraldehyde-3-phosphate dehydrogenase A OS=Escherichia coli (strain K12) OX | sp\|P0A9B2\|G3P1_ECOLI | 36 kDa | 10 |

| Identified Proteins (51) | Accession Number | Molecular Weight | SpC Sample 2 |
|---|---|---|---|
| Dps_TmT1\|SequenceB\|SequenceB | Custom\|SequenceB\|S | 32 kDa | 34 |
| Glutamine synthetase OS=Escherichia coli (strain K12) OX=83333 GN=glnA PE=1 SV=2 | sp\|P0A9C5\|GLN1B_E | 52 kDa | 13 |
| Elongation factor Tu 1 OS=Escherichia coli (strain K12) OX=83333 GN=tufA PE=1 SV=1 | sp\|P0CE47\|EFTU1_EC | 43 kDa | 9 |
| Biosynthetic arginine decarboxylase OS=Escherichia coli (strain K12) OX=83333 GN=speA | sp\|P21170\|SPEA_ECC | 74 kDa | 9 |
| Chaperonin GroEL OS=Escherichia coli (strain K12) OX=83333 GN=groEL PE=1 SV=2 | sp\|P0A6F5\|CH60_EC | 57 kDa | 7 |

| Identified Proteins (166) | Accession Number | Molecular Weight | SpC Sample 3 (Nat_MxT1) | Spc Sample 5 (Nat_MxT3) |
|---|---|---|---|---|
| Nat_MxT1/3\|SequenceC\|SequenceC | Custom\|SequenceC\| | 32 kDa | 253 | 224 |
| Ribonucleoside-diphosphate reductase 1 subunit alpha OS=Escherichia coli (strain K12) | sp\|P00452\|RIR1_ECC | 86 kDa | 0 | 2 |
| Maltodextrin phosphorylase OS=Escherichia coli (strain K12) OX=83333 GN=malP PE=1 | sp\|P00490\|PHSM_EC | 91 kDa | 0 | 6 |
| RNA polymerase sigma factor RpoD OS=Escherichia coli (strain K12) OX=83333 GN=rpoD | sp\|P00579\|RPOD_EC | 70 kDa | 0 | 8 |
| Beta-galactosidase OS=Escherichia coli (strain K12) OX=83333 GN=lacZ PE=1 SV=2 | sp\|P00722\|BGAL_EC | 116 kDa | 7 | 12 |

| Identified Proteins (67 | Accession Number | Molecular Weight | SpC Sample 4 (Dps_MxT1) | SpC Sample 6 (Dps_MxT3) |
|---|---|---|---|---|
| Dps_MxT1/3\|SequenceD\|SequenceD | Custom\|SequenceD\| | 33 kDa | 122 | 135 |
| 30S ribosomal protein S6 OS=Escherichia coli (strain K12) OX=83333 GN=rpsF PE=1 SV=1 | sp\|P02358\|RS6_ECOL | 16 kDa | 4 | 3 |
| 30S ribosomal protein S7 OS=Escherichia coli (strain K12) OX=83333 GN=rpsG PE=1 SV=3 | sp\|P02359\|RS7_ECOL | 20 kDa | 6 | 5 |
| 50S ribosomal protein L15 OS=Escherichia coli (strain K12) OX=83333 GN=rplO PE=1 SV=1 | sp\|P02413\|RL15_ECO | 15 kDa | 3 | 4 |
| Outer membrane porin F OS=Escherichia coli (strain K12) OX=83333 GN=ompF PE=1 SV=1 | sp\|P02931\|OMPF_EC | 39 kDa | 3 | 2 |

| Identified Proteins (170) | Accession Number | Molecular Weight | SpC Sample 7 |
|---|---|---|---|
| Nat_QtT4\|SequenceE\|SequenceE | Custom\|SequenceE | 32 kDa | 273 |
| Type II NADH:quinone oxidoreductase OS=Escherichia coli (strain K12) OX=83333 GN=ndh PE=1 S | sp\|P00393\|NDH_E | 47 kDa | 2 |
| Maltodextrin phosphorylase OS=Escherichia coli (strain K12) OX=83333 GN=malP PE=1 SV=7 | sp\|P00490\|PHSM_ | 91 kDa | 9 |
| Beta-galactosidase OS=Escherichia coli (strain K12) OX=83333 GN=lacZ PE=1 SV=2 | sp\|P00722\|BGAL_E | 116 kDa | 10 |
| Alanine--tRNA ligase OS=Escherichia coli (strain K12) OX=83333 GN=alaS PE=1 SV=2 | sp\|P00957\|SYA_EC | 96 kDa | 4 |

| Identified Proteins (113) | Accession Number | Molecular Weight | SpC Sample 8 |
|---|---|---|---|
| Dps_QtT4\|SequenceF\|SequenceF | Custom\|SequenceF\|S | 34 kDa | 263 |
| Maltodextrin phosphorylase OS=Escherichia coli (strain K12) OX=83333 GN=malP PE=1 SV=7 | sp\|P00490\|PHSM_EC | 91 kDa | 13 |
| RNA polymerase sigma factor RpoD OS=Escherichia coli (strain K12) OX=83333 GN=rpoD PE= | sp\|P00579\|RPOD_EC | 70 kDa | 5 |
| Beta-galactosidase OS=Escherichia coli (strain K12) OX=83333 GN=lacZ PE=1 SV=2 | sp\|P00722\|BGAL_EC | 116 kDa | 5 |
| 30S ribosomal protein S6 OS=Escherichia coli (strain K12) OX=83333 GN=rpsF PE=1 SV=1 | sp\|P02358\|RS6_ECOL | 16 kDa | 9 |

**Figure S2.** A) Native PAGE gel analysis of Nat_Encs and Dps_Encs stained with Coomassie Blue to visualize protein. Minor higher molecular weight bands for each Nat_Enc and Dps_Enc that were subjected to mass spectrometry identification are indicated by black boxes. B) Mass spectrometry results showing that all higher molecular weight bands represent the respective Nat_Encs or Dps_Encs. Only the top 5 most prevalent protein species are shown. Nat_Encs and Dps_Encs sequences are highlighted in gray.

| | Dps_TmT1 | Dps_MxT1 | Dps_MxT3 | Dps_QtT4 |
|---|---|---|---|---|
| Change in luminal charges upon Dps_N fusion | +180 | +180 | +540 | +720 |
| Change in luminal surface charge density upon Dps_N fusion($nm^{-2}$) | +0.177 | +0.398 | +0.254 | +0.177 |

**Figure S3.** Table showing the change in luminal charge and approximate luminal surface charge density upon Dps-N fusion in engineered Dps_Encs. Encapsulin shell thickness was assumed to be 3 nm. Interior surface area was calculated as follows: $4\pi r^2$ with r = shell radius - 3 nm, e.g., Tm: r = 12 nm - 3 nm = 9 nm. To obtain approximate luminal charge densities, we assumed the native luminal charge to be approximately neutral (or at least negligible, compared to the charge increase caused by Dps-N). Approximate luminal charge was calculated as follows: 3 x # of protomers / interior surface area.

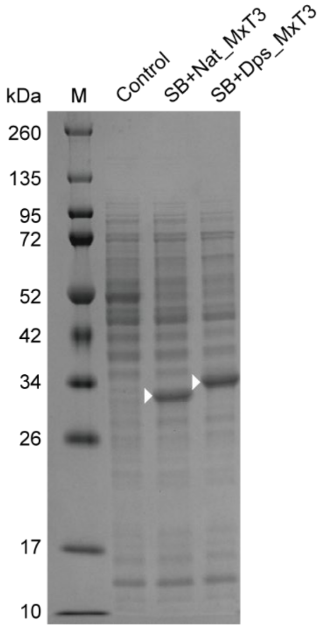


**Figure S4.** SDS-PAGE analysis of cell lysates from the control *E.coli* BL21(DE3) strain without any transformed plasmid, SB+Nat_MxT3, and SB+Dps_MxT3, showing that Nat_MxT3 and Dps_MxT3 are properly expressed at comparable levels. Bands corresponding to Nat_MxT3 and Dps_MxT3 are indicated by white arrows.

**Table S1.** Protein sequences of all proteins used in this study.

| Construct | Protein sequence |
|---|---|
| Nat_TmT1 | MEFLKRSFAPLTEKQWQEIDNRAREIFKTQLYGRKFVDVEGPYGWEYAAHPLGE<br>VEVLSDENEVVKWGLRKSLPLIELRATFTLDLWELDNLERGKPNVDLSSLEETVR<br>KVAEFEDEVIFRGCEKSGVKGLLSFEERKIECGSTPKDLLEAIVRALSIFSKDGIEG<br>PYTLVINTDRWINFLKEEAGHYPLEKRVEECLRGGKIITTPRIEDALVVSERGGDF<br>KLILGQDLSIGYEDREKDAVRLFITETFTFQVVNPEALILLKF |
| Nat_MxT1/3 | MPDFLGHAENPLREEEWARLNETVIQVARRSLVGRRILDIYGPLGAGVQTVPYD<br>EFQGVSPGAVDIVGEQETAMVFTDARKFKTIPIIYKDFLLHWRDIEAARTHNMPLD<br>VSAAAGAAALCAQQEDELIFYGDARLGYEGLMTANGRLTVPLGDWTSPGGGFQ<br>AIVEATRKLNEQGHFGPYAVVLSPRLYSQLHRIYEKTGVLEIETIRQLASDGVYQS<br>NRLRGESGVVVSTGRENMDLAVSMDMVAAYLGASRMNHPFRVLEALLLRIKHP<br>DAICTLEGAGATERR |
| Nat_QtT4 | MNKSQLYPDSPLTDQDFNQLDQTVIEAARRQLVGRRFIELYGPLGRGMQSVFND<br>IFMESHEAKMDFQGSFDTEVESSRRVNYTIPMLYKDFVLYWRDLEQSKALDIPID<br>FSVAANAARDVAFLEDQMIFHGSKEFDIPGLMNVKGRLTHLIGNWYESGNAFQDI<br>VEARNKLLEMNHNGPYALVLSPELYSLLHRVHKDTNVLEIEHVRELITAGVFQSPV<br>LKGKSGVIVNTGRNNLDLAISEDFETAYLGEEGMNHPFRVYETVVLRIKRPAAICT<br>LIDPEE |
| Dps_TmT1 | MSTAKLVKSKATNGGSGGSEFLKRSFAPLTEKQWQEIDNRAREIFKTQLYGRKFV<br>DVEGPYGWEYAAHPLGEVEVLSDENEVVKWGLRKSLPLIELRATFTLDLWELDN<br>LERGKPNVDLSSLEETVRKVAEFEDEVIFRGCEKSGVKGLLSFEERKIECGSTPK<br>DLLEAIVRALSIFSKDGIEGPYTLVINTDRWINFLKEEAGHYPLEKRVEECLRGGKII<br>TTPRIEDALVVSERGGDFKLILGQDLSIGYEDREKDAVRLFITETFTFQVVNPEALI<br>LLKF |
| Dps_MxT1/3 | MSTAKLVKSKATNGGSGGSPDFLGHAENPLREEEWARLNETVIQVARRSLVGRR<br>ILDIYGPLGAGVQTVPYDEFQGVSPGAVDIVGEQETAMVFTDARKFKTIPIIYKDFL<br>LHWRDIEAARTHNMPLDVSAAAGAAALCAQQEDELIFYGDARLGYEGLMTANG<br>RLTVPLGDWTSPGGGFQAIVEATRKLNEQGHFGPYAVVLSPRLYSQLHRIYEKTG<br>VLEIETIRQLASDGVYQSNRLRGESGVVVSTGRENMDLAVSMDMVAAYLGASRM<br>NHPFRVLEALLLRIKHPDAICTLEGAGATERR |
| Dps_QtT4 | MSTAKLVKSKATNGGSGGSNKSQLYPDSPLTDQDFNQLDQTVIEAARRQLVGRR<br>FIELYGPLGRGMQSVFNDIFMESHEAKMDFQGSFDTEVESSRRVNYTIPMLYKD<br>FVLYWRDLEQSKALDIPIDFSVAANAARDVAFLEDQMIFHGSKEFDIPGLMNVKG<br>RLTHLIGNWYESGNAFQDIVEARNKLLEMNHNGPYALVLSPELYSLLHRVHKDTN<br>VLEIEHVRELITAGVFQSPVLKGKSGVIVNTGRNNLDLAISEDFETAYLGEEGMNH<br>PFRVYETVVLRIKRPAAICTLIDPEE |
| eGFP_MxTP | MVSKGEELFTGVVPILVELDGDVNGHKFSVSGEGEGDATYGKLTLKFICTTGKLP<br>VPWPTLVTTLTYGVQCFSRYPDHMKQHDFFKSAMPEGYVQERTIFFKDDGNYK<br>TRAEVKFEGDTLVNRIELKGIDFKEDGNILGHKLEYNYNSHNVYIMADKQKNGIKV<br>NFKIRHNIEDGSVQLADHYQQNTPIGDGPVLLPDNHYLSTQSALSKDPNEKRDH<br>MVLLEFVTAAGITLGMDELYKGGSGGSPEKRLTVGSLRR |
